# The impact of the newly licensed dengue vaccine in endemic countries

**DOI:** 10.1101/074062

**Authors:** M. Aguiar, Nico Stollenwerk, Scott B. Halstead

## Abstract

**Background:** With approximately 3 billion people at risk of acquiring the infection, dengue fever is now considered the most important mosquito-borne viral disease in the world, with 390 million dengue infections occurring every year, of which 96 million manifest symptoms with any level of disease severity. Treatment of uncomplicated dengue cases is only supportive and severe dengue cases require hospital intensive care. A vaccine now licensed in several countries and developed by Sanofi Pasteur (CYD-TDV, named Dengvaxia), is able to protect, in the first 25 months of the two Phase III, 66% of a subset of 9 − 16 year old participants. However, a significantly lower efficacy (including negative vaccine efficacy) was noted for children younger than 9 years of age.

**Methodology/Principal Findings:** Analysis of year 3 results of phase III trials of Dengvaxia suggest high rates of protection of vaccinated partial dengue immunes but high rates of hospitalizations during breakthrough dengue infections of persons who were vaccinated when seronegative, with vaccine appearing to induce enhancing antibodies (ADE). An age structured model was developed based on Sanofi’s recommendation to vaccinate persons age 945 years in dengue endemic countries. The model was used to explore the clinical burden of two vaccination strategies: 1) Vaccinate 4 or 20% of individuals, ages 9 − 45 years, seropositives and seronegatives, and 2) vaccinate 4 or 20% of individuals, ages 9 − 45 years, who are dengue immune only.

**Conclusions/Significance:** Our results show that vaccinating dengue monotypic immune individuals prevents dengue hospitalizations, but at the same time dengue infections of vaccine-sensitized persons increases hospitalizations. When the vaccine is given only to partial immune individuals, after immunological screening of the population, disease burden decreases considerably.

**Author Summary:** Caused by four antigenically related but distinct serotypes a tetravalent vaccine is needed to protect against the huge burden of dengue disease. Dengvaxia is a vaccine candidate now licensed in several countries for individuals 9 − 45 years of age living in endemic countries with least 70% of seroprevalence. Modelers from Sanofi Pasteur have predicted that this vaccine has the potential to reduce by about 50% the disease burden within 5 years when 20% of an endemic country population is vaccinated, thus achieving a World Health Organization dengue prevention goal.

In this paper, mathematical modeling is used to investigate the impact of the newly licensed dengue vaccine using different scenarios. Our results show that to achieve significant reduction in disease burden, the vaccination program is most effective if it includes only individuals that have been already exposed to at least one dengue virus. Immunological screening of the population prior to vaccination is advised and vaccination strategies must be planned based on epidemiological disease dynamics for each specific endemic region.

## 1 Introduction

Epidemiological models have been important in understanding the spread of infectious diseases and to evaluate intervention strategies like vector control and vaccination. Mathematical models were introduced into infectious disease epidemiology in the early 20th century, and a series of deterministic compartmental models, such as e.g. the SIR (susceptible-infected-recovered) type model, have been proposed based on the flow patterns between compartments of hosts. Recently, most models try to incorporate several different aspects of the disease, including the duration of disease, duration of infectivity, infection rate, waning immunity, and so forth, bringing rich dynamic behavior in most simple models.

The dynamics of dengue disease and transmission reveals large fluctuations of disease incidence challenging mathematical models to explain the irregular behaviour of dengue epidemics. Dengue fever (DF) is caused by four antigenically related but distinct serotypes (DENV-1 to DENV-4). Infection by one serotype confers life-long immunity to that serotype and a period of temporary cross-immunity (TCI) to other serotypes. The clinical response on exposure to a second serotype is complex and may depend on factors such as patient age, dengue type or strain, sequence of infection and the interval between infection by one serotype and exposure to a second serotype. Epidemiological studies support the association of severe disease (dengue haemorrhagic fever (DHF) and dengue shock syndrome (DSS)) with secondary dengue infection. There is good evidence that sequential infection increases the risk of developing DHF/DSS [1–6], due to a process described as antibody-dependent enhancement (ADE), where the pre-existing antibodies to previous dengue infection do not neutralize but rather enhance the new infection.

Early models have described multi-strain interactions leading to complex behaviour via ADE, e.g.[7–9], but always neglecting the effect of temporary cross-immunity. Then, incorporation of TCI in complex models tested the impact of ADE acting to increase the infectivity of secondary infections [10–12]. More recently, complex dynamics (known as deterministic chaos) was found in wider and more biologically realistic parameter region able to describe disease epidemiology, no longer needing to assume infectivity of secondary infections as much greater than the infectivity of primary infections [13, 14]. The minimalistic model by Aguiar et al. [14] successfully described large fluctuations observed in empirical outbreak data [14–16]. It was estimated that secondary dengue infections contribute to transmission at a rate that is at least 10% less than primary infections, anticipating results published recently in [17], demonstrating that persons with inapparent DENV infections were more infectious to mosquitoes than clinically symptomatic patients. As described in [18], more than the detailed number of strains, a combination of TCI and ADE are the most important features driving the complex dynamics observed in dengue fever epidemiology.

Treatment of uncomplicated dengue cases is only supportive, and severe dengue cases require hospitalization. A vaccine is needed to protect against the huge burden of dengue disease resulting in a requirement that a protective immune response be mounted to all four serotypes [19]. The dengue vaccine candidate available for licensing is a live attenuated tetravalent dengue vaccine developed by Sanofi Pasteur (CYD-TDV, named Dengvaxia). Phase III trials were successfully completed in the Asian-Pacific region and in Latin American countries and vaccine efficacy estimates for confirmed dengue cases, with wide error margins, were recently published for each trial [20, 21]. According to the manufacturer the vacine efficacy has reduced dengue disease prevalence, during the first 25 months of Phase III, by 66% of a subset of 9 − 16 year olds [22].

In November 2015, modellers from Sanofi Pasteur have predicted that Dengvaxia has the potential to reduce the dengue disease burden by about 50% within 5 years, when 20% of an endemic country’s population is vaccinated [23], thus achieving a World Health Organization (WHO) dengue prevention goal. However, analysis of year 3 results of phase III trials suggest high rates of protection of vaccinated partial dengue immunes, but also high rates of hospitalizations during breakthrough dengue infections of persons who were vaccinated when seronegative [24]. The latter result indicates that Dengvaxia appears to induce dengue infection-enhancing antibodies (ADE) [25].

Sanofi recommends giving vaccine to persons age 9 − 45 years in dengue endemic countries and if this strategy is deployed, dengue infections of monotypic dengue immunes will be prevented, but at the same time, “a theoretical risk” of dengue infections of vaccine-sensitized persons resulting in hospitalizations is expected [27]. A significant demand for the vaccine is expected, estimated at 60 million to 80 million doses per year [30]. To date, Mexico, the Philippines, Brazil, El Salvador, Costa Rica and Paraguay have licensed DengVaxia for use in their countries [31–36]. One million doses of the vaccine were shipped to the Philippines where the government has implemented the Dengue School-Based Immunization program that commenced on April 4 [37]. On April 15, 2016, the WHO Strategic Advisory Group of Experts (SAGE) on immunisation recommended Dengvaxia be used in regions with high endemicity, as indicated by seroprevalence of approximately 70% or greater in the targeted age group [26–28], when modeled reduced dengue hospitalizations by 10 − 30% over 30 years. Although the vaccine was considered safe, one can read in the WHO-SAGE report that “individuals who were seronegative at the time of vaccination were predicted to be at increased risk for hospitalization.” It was also mentioned that “In travellers unlikely to have already had dengue, vaccination may be substantially less beneficial (and there is a theoretical risk that it may be harmful), analogous to seronegative individuals living in endemic settings” [27].

In the present manuscript, using a Bayesian approach [38], we present the statistical description of the vaccine efficacy estimates performed for the Sanofi Pasteur tetravalent dengue vaccine trials data. Vaccine efficacy for preventing hospitalization of virologically confirmed dengue cases is estimated, based on relative risk information publicly available in [24, 27]. Using the published mathematical modeling approach [13–15, 18], an age structured model is developed based on Sanofi’s recommendation to vaccinate persons age 9 − 45 years in dengue endemic countries. Here we explore the clinical outcome of two vaccination strategies: 1) Vaccinate 4 or 20% of individuals, ages 9 − 45 years, seropositives and seronegatives, and 2) vaccinate 4 or 20% of individuals, ages 9 − 45 years, who are dengue immune only. We test for the assumed protective efficacy vaccine given in the province of Chiang Mai in Thailand. The most effective vaccine implementation scenario is identified and the impact of Dengvaxia administration is discussed.

## Methods

### 1.1 Modelling the efficacy of the Sanofi Pasteur dengue vaccine

Sanofi Pasteur’s vaccine trials were conducted in countries in the Asian-Pacific region and in five Latin American countries, with more than 35.000 children receiving either vaccine or placebo [20, 21]. Based on the number of virologically confirmed dengue cases after the third injection, the vaccine efficacy was estimated and reported separately for each of the study regions.

For modelling this vaccine trial, where there is one data point for the control (placebo) group and one for the vaccine group receiving respectively three doses of placebo or the tetravalent vaccine, we have employed a simple epidemiological susceptible *S* and infected *I* linear infection model with constant force of infection [39]. With the total number of participants in the control group *N*_*c*_ = *S*_*c*_ + *I*_*c*_, we have the scheme

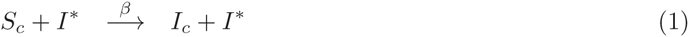

where susceptible individuals *S*_*c*_ become infected *I*_*c*_ with a natural infection rate *β* after a time interval *T*:= *t* − *t*_0_. Here *T* = 12 months for the actual vaccine trial. The underlying model hypothesis is that infection can be acquired from outside the considered control group when meeting an external infected individual *I*∗. For the vaccine group we have the same epidemiological process, but expect reduced infectivity *c* ⋅ *β* (when *c <* 1) instead of *β*. The linear infection model for the vaccine group is represented by the scheme below

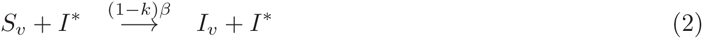

where *S*_*v*_ + *I*_*v*_ = *N*_*v*_ and the vaccine efficacy *k* can be written explicitly as *k*:= 1 − *c*.

#### 1.1.1 Vaccine efficacy for confirmed dengue cases: years 1-2

Using the data given by the Sanofi-Pasteur dengue vaccine trials in the Asian-Pacific region (CYD14) as reported in [20] and the Latin American countries (CYD15) as reported in [21], we could estimate the overall vaccine efficacy for virologically confirmed dengue cases via the Bayesian approach [39–41] to obtain a probability *p*(*k*|*I*_*v*_, *I*_*c*_) for the vaccine efficacy *k*) with infected individuals *I*_*v*_ in the vaccine group and *I*_*c*_ in the control group. For more detailed calculations, see [38].

We obtained a statistical description of the vaccine trial data that were in very good agreement with the results for the overall vaccine efficacy for confirmed dengue cases (56.5% with 95%-CI (43.8; 66.4) published in [20] versus our results of 56.56% with 95%-CI (44.4; 66.2) and 60.8% with 95%-CI (52.0; 68.0) published in [21] versus our results of 60.76% with 95%-CI (52.2; 70.5)) for the Asian-Pacific Region, shown in Fig.1 (a) using dark orange curve, and for the Latin America countries, shown in Fig.1 (a) using light orange curve. Note that the uninformed Bayesian prior might have a slight influence mainly in the confidence interval values.

**Figure 1.**
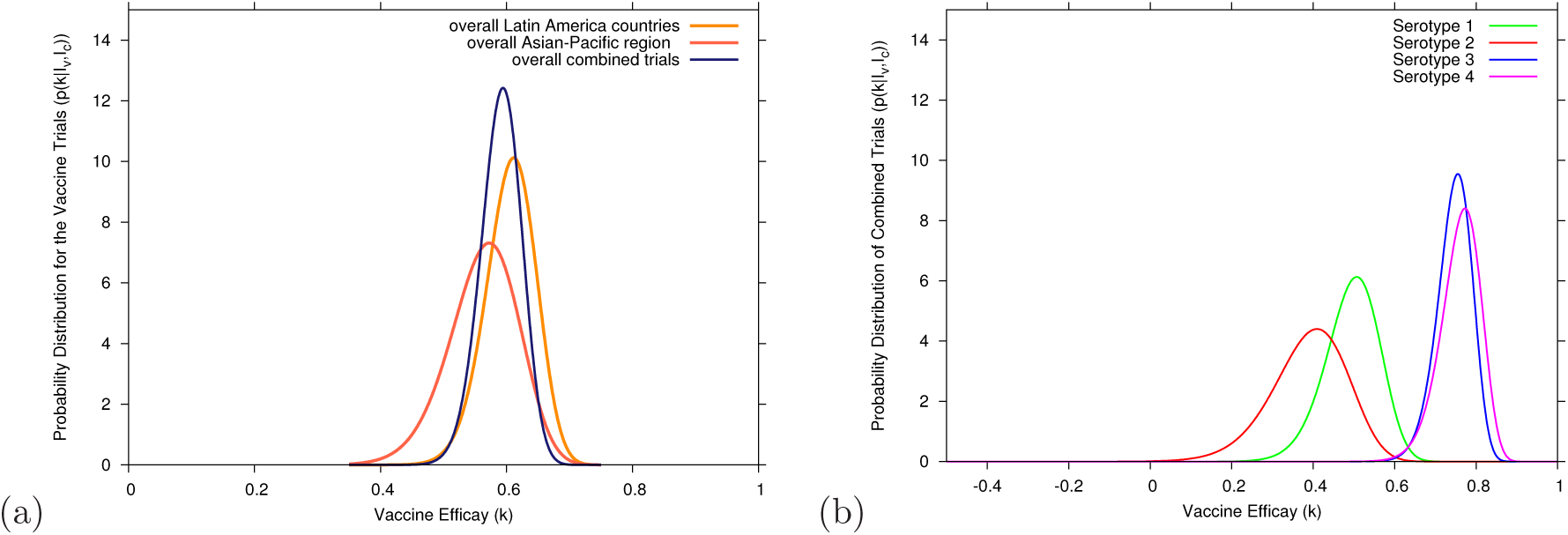
Bayesian efficacy analysis for Sanofi Pasteur’s tetravalent dengue vaccine trials. In (a) the Bayesian estimate of the overall vaccine efficacy for confirmed dengue cases. The dark orange curve shows the estimates for the Asian-Pacific region, the light orange curve shows the estimates for the Latin America countries and the dark blue curve shows the estimates for the combined Asian-Pacific and Latin America studies. In (b) the Bayesian estimate of the serotype specific vaccine efficacy; Serotypes 1, 2, 3 and 4 are shown respectively in green, red, blue and magenta.

Both trials were carried out using the same protocol designed to smoothen differences in population diversity and the climatic conditions. Knowing that larger sample size will provide a more accurate parameter estimation and increase the chance of statistical significance, we estimated global vaccine efficacy assuming that the variety in trial scenario among the Asian-Pacific region has the same degree of diversity as in the trial scenario among the Latin American countries, and that diversities observed between Asian-Pacific and the Latin American countries are comparable.

Using the combined data from Sanofi-Pasteurs dengue vaccine trials (CDY14+CYD15) we estimated the overall vaccine efficacy, via the Bayesian approach, to obtain a probability for the combined vaccine efficacy based on the empirical data from Asian-Pacific and the Latin American trials [20, 21].

The Bayesian estimate of the combined vaccine efficacy trials is *k* = 59.2% with a 95%-CI of (52.4; 65.0) is shown in Fig.1 (a), as dark blue curve. It is slightly below the 60.8% obtained in the Latin American trial, but higher than the 56.6% obtained in the Asian-Pacific trial. For the serotype specific vaccine efficacy, shown in Fig.1 (b), a tendency for a grouped efficacy is observed, where serotypes *DEN*_3_ and *DEN*_4_ are grouped together with a relatively good efficacy (approx. 78%) and serotype *DEN*_1_ (approx. 50%) and serotype *DEN*_2_ (approx. 40%) appear with an intermediate efficacy. Any possible common efficacy for all serotypes is statistical excluded [42].

#### 1.1.2 Vaccine efficacy for hospitalized dengue cases: year 3

Results from years 1−2 demonstrated an intermediate efficacy against a wide spectrum of disease observed during the surveillance phase. Vaccine efficacy for seronegative individuals at baseline (individuals that have never been infected by a dengue virus prior to the vaccine trial) was found to be considerably smaller than the efficacy for individuals who were seropositive at baseline. During the stage where cases were monitored in hospitals, during year 3 there was a higher incidence of hospitalization among children ages 5 years and younger (in total, 20 children (1.4%) being hospitalized in the vaccine group and only 2 (0.1%) hospitalizations observed in the control group) [24].

The relative risk of hospitalization for children of all ages in 11 trial sites was estimated to be about 1 (1.04, with the 95%-CI to be of (90.52; 2.19)). For children younger than 9 years of age and specifically for children between 2 − 5 years of age, the relative risk was estimated to be 1.58 (95%-CI (0.61; 4.83)) and 7.45 (95%-CI (1.15; 313.8)) respectively [24].

Similar to the analysis performed in section 1.1.1, we now use the data for the annual incidence of dengue cases hospitalization for the Asian-Pacific region (CYD14) and for the Latin American countries (CYD15) presented as relative risk in [24], to estimate the vaccine efficacy for hospitalized dengue cases.

**Figure 2.**
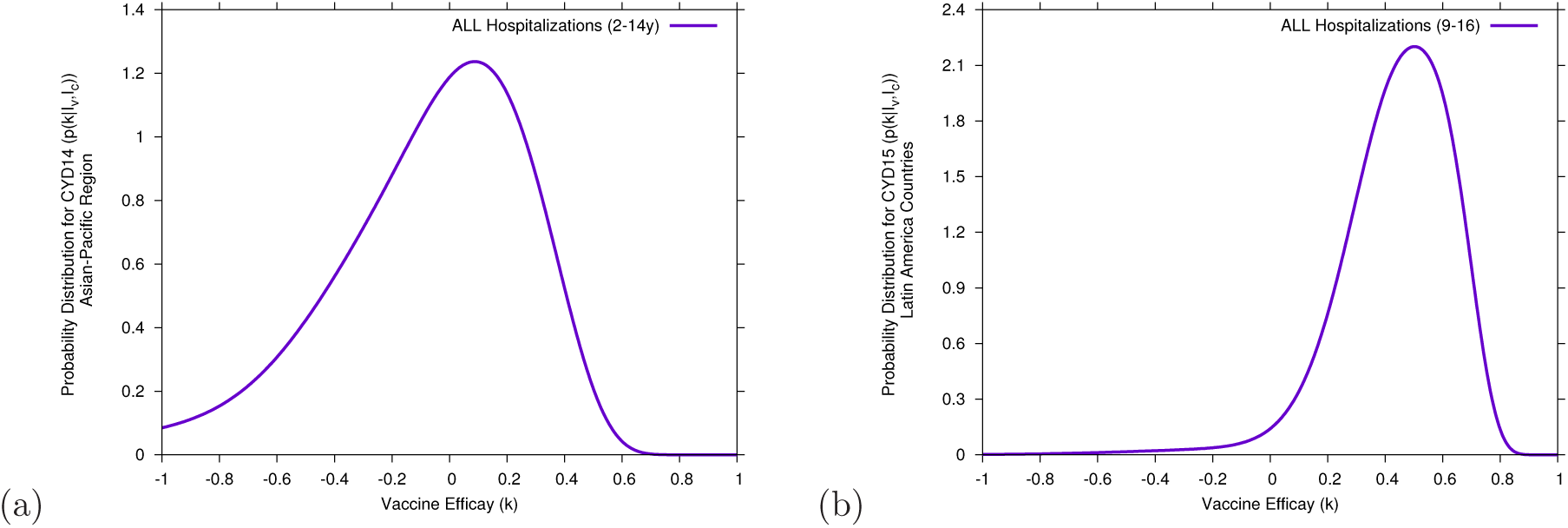
Bayesian estimate of the vaccine efficacy for dengue hospitalized cases. In (a) for the Asian-Pacific region (CYD14 trial) and in (b) for the Latin American countries (CYD15 trial).

The Bayesian estimate of the vaccine efficacy for all hospitalized cases (2-14 years old) in the CYD14 trial is *k* = −3.2% with a 95%-CI of [−108.8; 45.4], shown in Fig.2 (a), and for all hospitalized cases (9-16 years old) in the CYD15 was found to be *k* = 46.0% with a 95%-CI of [−0.6; 72.2], shown in Fig.2 (b). The observed difference in vaccine efficacy estimation per trial is explained since children below 9 years of age, who are expected to be mostly seronegatives, were not recruited in the CYD15 trial.

A negative vaccine efficacy was estimated in the CYD14 trial, with vaccine disease-enhancement in younger children. In Fig.3 (a), the Bayesian estimate of the vaccine efficacy for hospitalized cases in children under 9 years of age is *k* = −53.6% with a 95%-CI of (−279.8; 38.2), whereas for children between 2 − 5 years of age, shown Fig.3 (b), the Bayesian estimate of the vaccine efficacy for hospitalized cases is *k* = −530.6% with a 95%-CI of (−631.6; −401.4). Here, any positive vaccine efficacy is statistically rejected.

**Figure 3.**
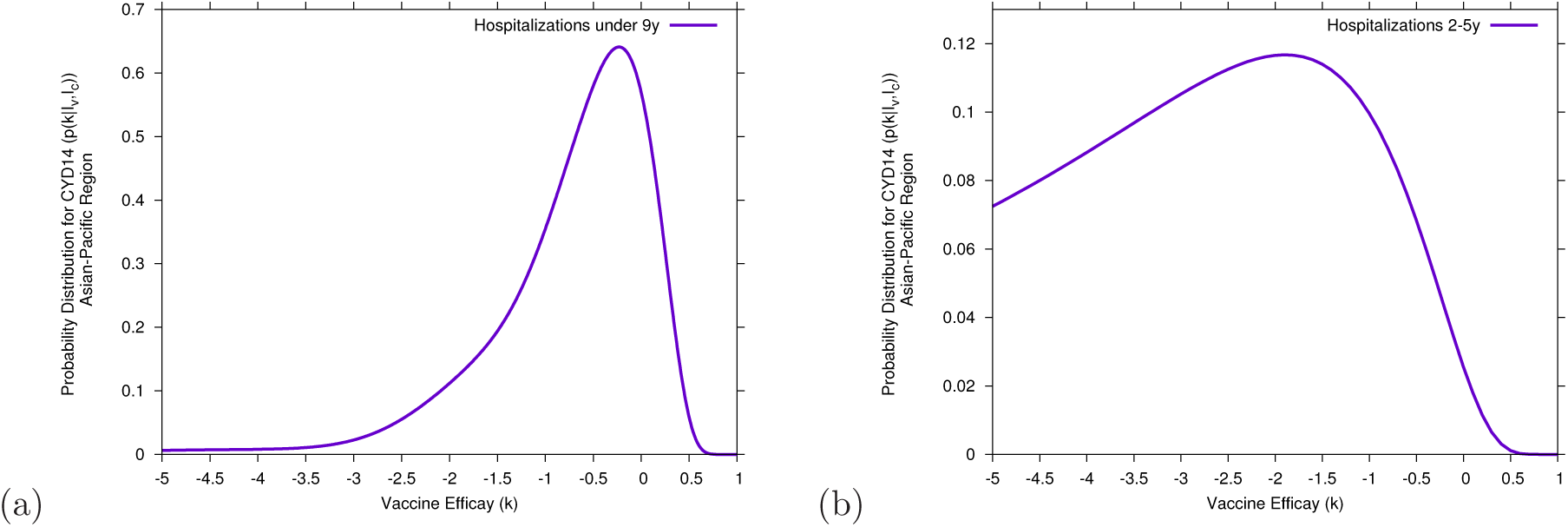
Bayesian estimate of the CYD14 vaccine efficacy for hospitalized dengue cases in children unde 9 years of age. In (a) all individuals under 9 years of age and in (b) children between 2 − 5 years of age.

#### 1.1.3 Vaccine efficacy for cumulative hospitalized dengue cases: years 3-5 for CYD14 and years 3-6 for CYD23/57

At the time of first submission of this article, on April 12 2016, only data for hospitalized dengue cases during year 3 of CYD14 trial were available [24]. Analysis using the hospitalized dengue cases during the years 4 and 5 of CYD14 trial can now be performed. The updated data were obtained from Table 8 presented in the WHO-SAGE background paper [27].

**Figure 4.**
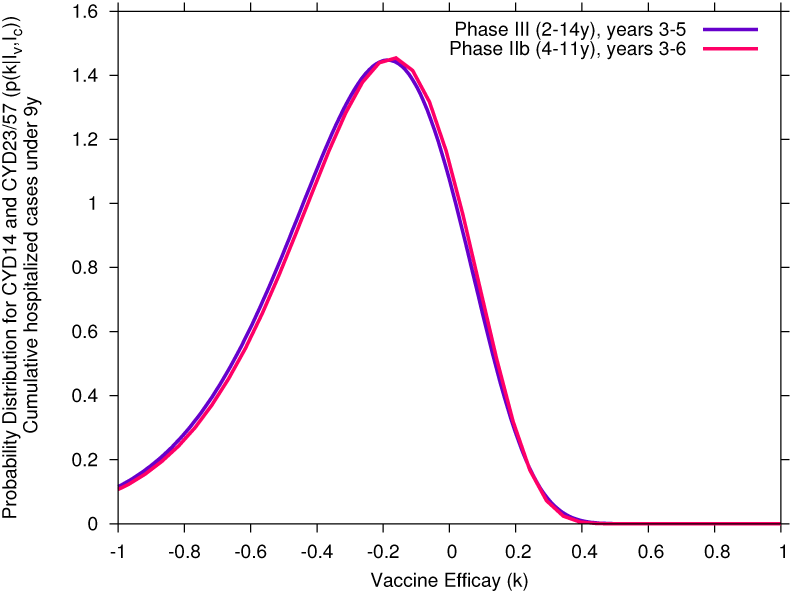
Bayesian estimate of vaccine efficacy for cumulative hospitalized dengue cases. In (a) individuals under 9 years of age during years 3-5 in the CYD14 trial and in (b) individuals under 9 years of age during years 1-6 in the CYD23/57 trial.

Using the cumulative data for hospitalized cases of individuals under 9 years, during year 3-5 of the CYD14 trial, the Bayesian estimate of the vaccine efficacy was found to be *k* = −27.1% with a 95%-CI of [−99.0; 17.5]. For hospitalized cases of individuals under 9 years, during year 3-6 of the CYD23/57 trial, the Bayesian estimate of the vaccine efficacy was found to be similarly negative, with *k* = −26.2% with a 95%-CI of [−98.0; 0.14]. Note that vaccine efficacy estimation for the year 3 of the CYD14 trial (−53.6%) remains within the confidence interval described here.

These estimations, shown in Fig. 4, support the finding of Sanofi Pasteur’s vaccine-induced immune response sensitizing seronegative individuals to enhanced disease accompanying breakthrough DENV infections [25] of disease-enhancement observed in vaccinated seronegative individuals.

In the following, an age structured model is developed, based on Sanofi’s recommendation for vaccine administration to individuals 9 − 45 years of age. With the purpose of describing differences in vaccine efficacy in relation to the immunological status of individuals prior to vaccination. We define seropositive individuals to be those infected with at least one lifetime dengue infection and seronegative individuals, those never been infected with any dengue virus.

We agree with the manufacturer’s explanation for the observed increase in hospitalization in children under 9 years of age but believe the same phenomenon prevails for all age groups: “The risk of developing severe disease is higher for individuals with a secondary heterotypic infection than for those with a primary infection. This may be mimicked by CYD-TDV vaccination of seronegative individuals, whereby vaccination represents a primary-like infection dominated by one or a few serotypes, and diminishing responses lead to only short-term cross-protection. As cross-protection wanes (potentially rapidly, owing to low antibody titres), so vaccine efficacy is reduced, as discussed above. Furthermore, vaccinated individuals could be at greater risk of developing a severe or symptomatic secondary-like infection the first time they contract DENV: the vaccine could act as their primary infection, and the subsequent true primary wild-type DENV infection (which would otherwise be typically less severe) could simulate a secondary wild-type infection (which is typically more severe).” [29].

In our model, individuals 9-45 years of age are vaccinated. The population consists of a mixture of seronegative and seropositive individuals. In this model, unvaccinated individuals are assumed to develop a mild or asymptomatic disease outcome during a primary natural infection whereas secondary infection is often symptomatic, admitted to hospitals and classified as DF, DHF or DSS [16]. Here, vaccine efficacy for seropositive individuals is assumed to be close to 100%, preventing dengue infections and hospitalization of monotypic dengue immune individuals. Note that among trial participants 9 years of age or older, seropositivity was approximately 80%. However, vaccine efficacy in seronegative individuals of all ages are assumed to be the same as the vaccine efficacy described for hospitalized children younger than 9 years, since this age group had a higher proportion of seronegatives [27].

### 1.2 Mathematical modeling of future vaccine impact

We extend the minimalistic two-strain dengue model investigated previously by Aguiar et al. [13,14] which is able to capture differences between primary and secondary dengue infection, including temporary cross-immunity an important parameter for modeling dengue fever epidemiology [18]. We consider only two possible infections, primary and secondary, an assumption validated because of the low frequency of tertiary and quaternary infections among hospital cohorts [43].

The model is represented in Fig.5 using a state flow diagram.

**Figure 5.**
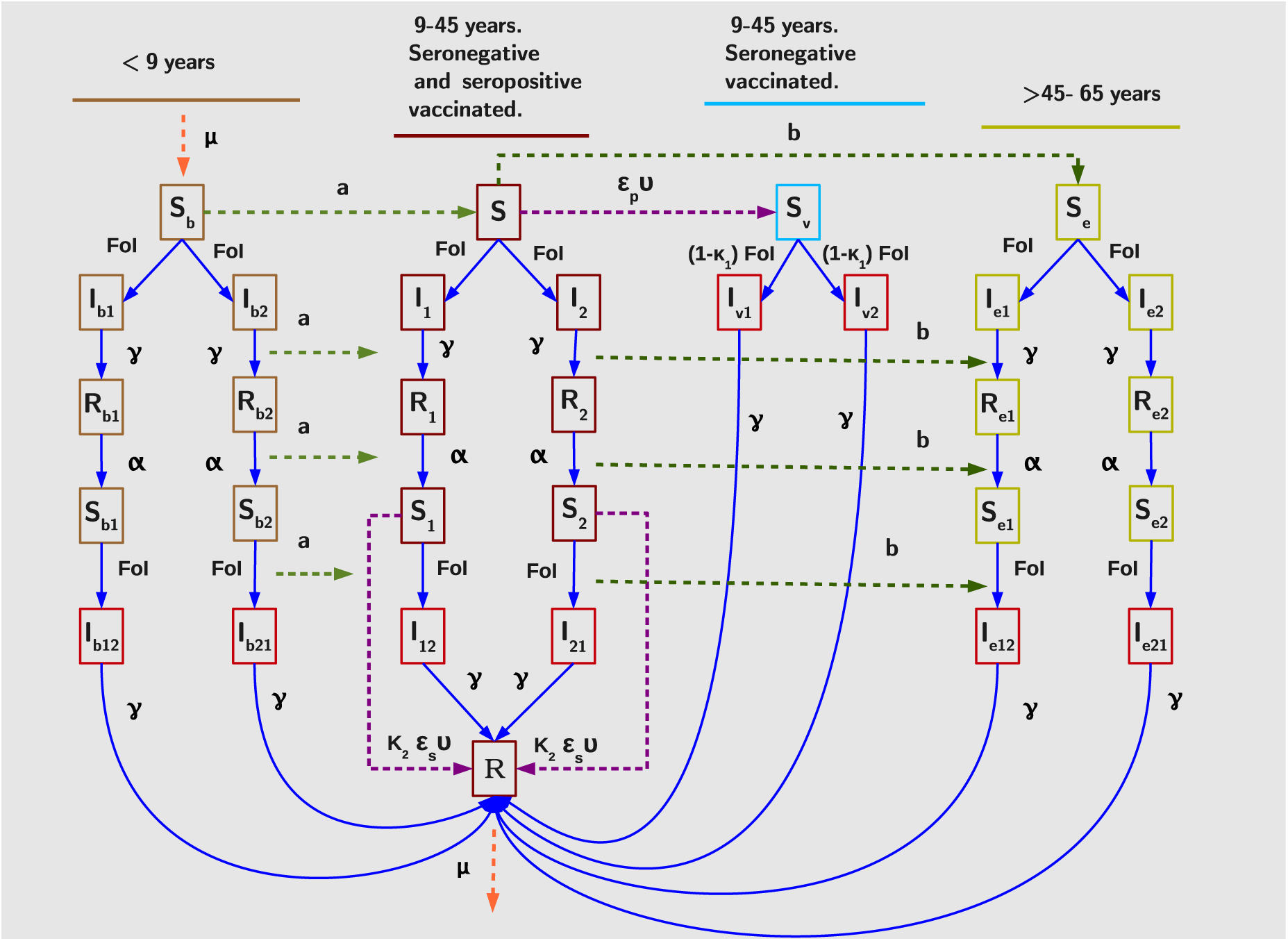
State flow diagram for age-structured two-strain dengue model with vaccination. Hospitalizations are only possible in the classes marked in red. Force of infection (FoI) is explicitly given by. *β*(*I*_1_ + *I*_2_ + *I*_*b*1_ + *I*_*b*2_ + *I*_*e*1_ + *I*_*e*2_ + *ρN* + *φ*(*I*_12_ + *I*_21_ + *I*_*b*12_ + *I*_*b*21_ + *I*_*e*12_ + *I*_*e*21)_ + *φv*(*I*_*v*1_ + *I*_*v*2_)).

The complete system of ordinary differential equations for the dengue vaccine epidemiological model is given by

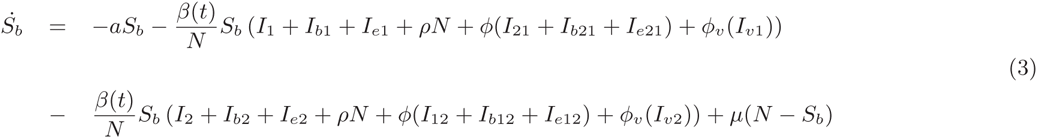

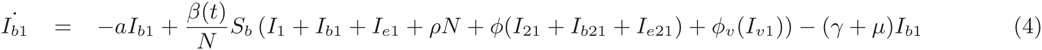

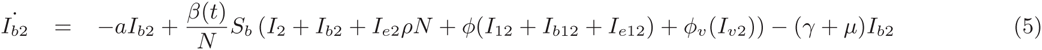

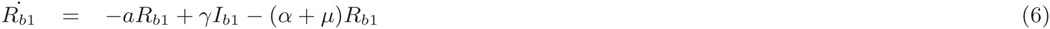

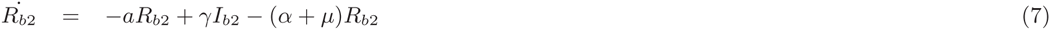

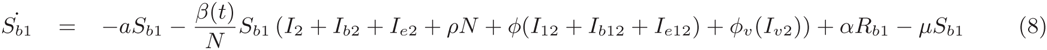

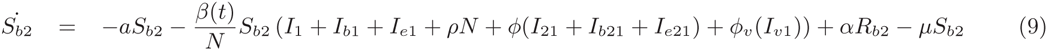

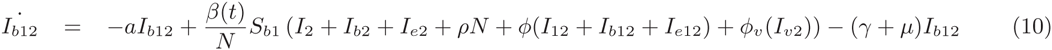

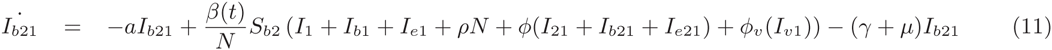

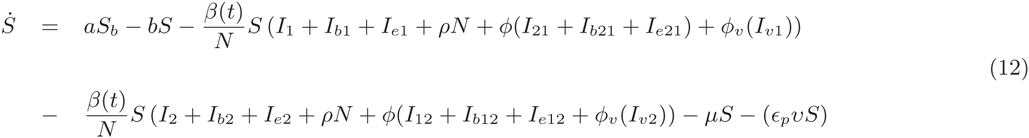

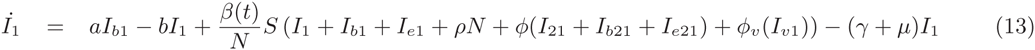

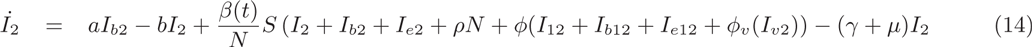

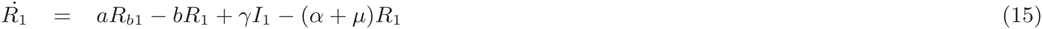

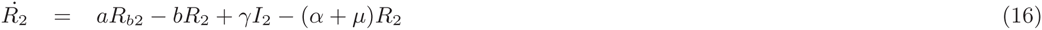

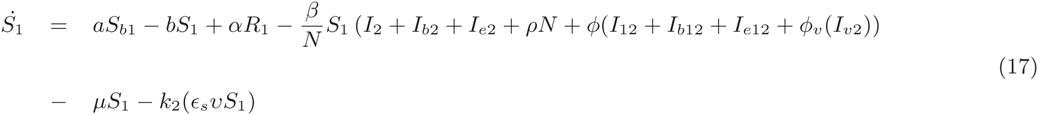

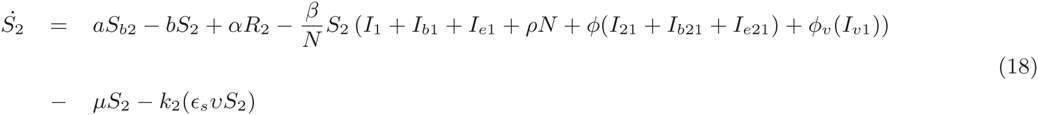

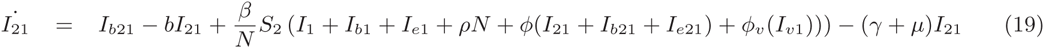

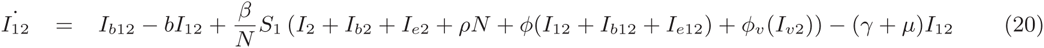

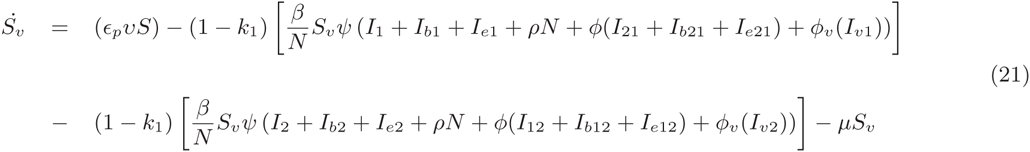

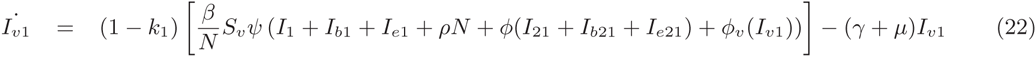

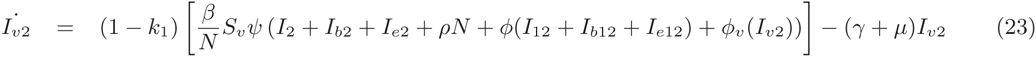

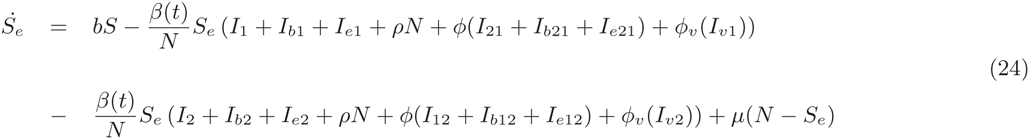

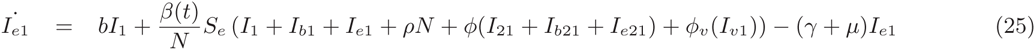

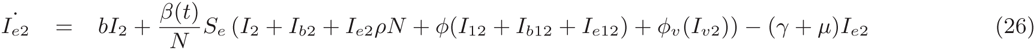

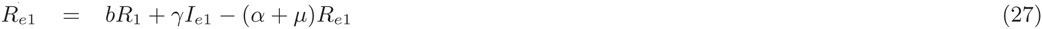

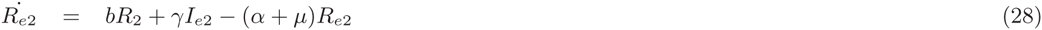

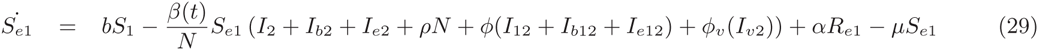

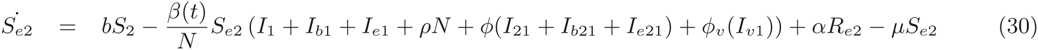

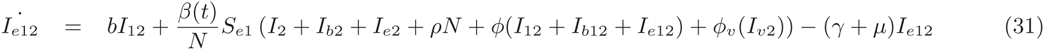

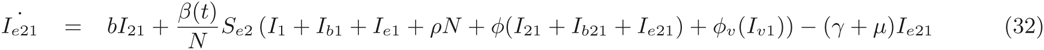

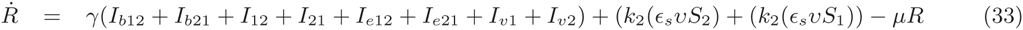

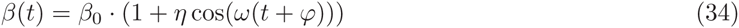

The parameter *β* takes the seasonal forcing into account as a cosine function and is given explicitly by where *β*_0_ the average infection rate, *η* is the degree of seasonality and *ϕ* is the phase of seasonality. The temporary cross-immunity period and the recovery rate are parametrized by *α* and *γ* respectively. The population size *N* is constant and the demography (birth and death rates) is parametrized by *µ*.

In this model, susceptible individuals can become infected also by meeting an infected individual from an external population, hence ((*β/N*) ⋅ *S* ⋅ *I* goes to (*β/N*) ⋅ *S* ⋅ (*I* + *ρ* ⋅ *N*)), contributing to the force of infection with an import parameter *ρ*. The parameters *φ* and *φ*_*v*_ are respectively the ratio of infection contribution to the force of infection from non-vaccinated seropositive individuals and vaccinated seronegative (vaccine-sensitized) individuals.

The effects of the vector dynamics are only taken into account by the force of infection parameters in the SIR-type model, but not modeling this mechanisms explicitly [44]. Like this, three different transmission rates are possible, depending of the immunological status of individuals transmitting the disease. Non-vaccinated individuals with a primary infection (*I*_1_ + *I*_2_ + *I*_*b*1_ + *I*_*b*2_ + *I*_*e*1_ + *I*_*e*2_) and with a secondary infection (*I*_12_ + *I*_21_ + *I*_*b*12_ + *I*_*b*21_ + *I*_*e*12_ + *I*_*e*21_) transmit disease with infection rate *β* and *φβ* respectively. Vaccinated seropositive individuals do not become infected and therefore do not transmit the disease, however, infected seronegative which were vaccinated (*I*_*v*1_ + *I*_*v*2_) will transmit the disease with infection rate *φ*_*v*_*β*.

An analysis of breakthrough hospitalizations for natural dengue infections in vaccinated seronegative children revealed an increased risk compared with hospitalizations accompanying secondary dengue infections [25]. Accordingly, a vaccine disease-enhancement factor *ψ* is introduced that conservatively increases risk of hospitalization of seronegative individuals acquiring a natural dengue infection by 10% (*S*_*v*_*ψ* going to *I*_*v*1_ or *I*_*v*2_).

The two-infection dengue model with serotype structure is constructed assuming three age classes: Individuals younger than 9 years (labeled with subscript *b*), individuals 9−45 years and individuals older than 45 years (labeled with subscript *e*) of age. Individuals younger than 9 and older than 45 years of age (see Fig.5, in light brown and dark yellow respectively) constitute the age classes that can not be vaccinated whereas individuals between 9 − 45 years of age (see Fig.5, in dark red) comprises the target population eligible to receive the vaccine. We recall that the population consists in a mixture of seronegative and seropositive individuals.

The dynamics for the two-infection dengue model is described as follows:

All individuals of every age start as seronegative susceptibles *S*_*b*_, *S* and *S*_*e*_ respectively and can acquire a natural first infection *I*_*b*1_ or *I*_*b*2_, *I*_1_ or *I*_2_ and *Ie*1 or *Ie*2 with a specific DENV-type 1 or 2. Dengue transmission occurs between susceptibles and individuals experiencing a primary or natural secondary dengue infection or a primary infection of a vaccinated seronegative individual. Note that the strain labeling 1 and 2 becomes important for secondary infections that can only occur with a different virus from the one causing the primary infection. Similarly to the assumptions in [26], individuals experiencing a secondary infection are assumed to have higher risk (than primarily infected individuals) of developing clinically apparent and severe disease with hospitalization.

Individuals recover from a primary infection *R*_*b*1_ or *R*_*b*2_, *R*_1_ or *R*_2_ and *R*_*e*1_ and *R*_*e*2_ with a recovery rate *γ* and after a period temporary cross-immunity *α*, become seropositive susceptibles *S*_*b*1_ or *S*_*b*2_, *S*_1_ or *S*_2_ and *S*_*e*1_ or *S*_*e*2_, now immune to one DENV type and able to acquire a natural secondary dengue infection *I*_*b*12_ or *I*_*b*21_, *I*_12_ or *I*_21_ and *I*_*e*12_ or *I*_*e*21_, with a different DENV type, respectively. Finally, individuals recover from a secondary infection *R*. Note that up to two dengue infections are possible before individuals reach the age of 9. When individuals reach age 9 years they move, with a transition rate *a* = 1/9 year (see light green arrows in the flow diagram, Fig. 5), to the compartments of disease related stages for individuals that are 9 − 45 years of age.

By vaccinating seronegative susceptible individuals *S*, a new class *S*_*v*_ is created (see light blue box in the flow diagram, Fig.5). The vaccine mimics the immunological effect of a primary dengue infection sensitizing them to a more severe secondary infection as has been suggested in [29]. As presented above, these sensitized individuals when naturally infected for the first time *I*_*v*1_ or *I*_*v*2_ have a 10% higher probability of hospitalization as compared to individuals acquiring the natural secondary dengue infection (*I*_12_ and *I*_21_). Thus, when seropositive individuals *S*_1_ and *S*_2_ are vaccinated the result is protection (going to *R*) while vaccinated seronegative individuals *S*_*v*_ becoming infected for the first time *I*_*v*1_ and *I*_*v*2_ have higher chance to develop severe disease and being hospitalized.

Individuals older than 45 years transition to a group that no longer receives vaccine with a transition rate *b* = 1/45 year (see dark green arrows in the flow diagram, Fig.5). In our model, vaccine is imple-mented such that the vaccination period is 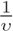, where eligible individuals receive 3 doses over a period of 1 year (see purple arrows in the flow diagram, Fig.5). The vaccination coverage per year is *ǫ* in an endemic country among individuals 9 to 45 years old, seronegative or seropositive prior to vaccination.

The SAGE committee advised giving vaccine in populations composed of 70% of seropositive individuals. It should be noted that the remaining 30% of the population are seronegative. Published models suggest that when this 30% is included in vaccination campaigns an overall disease burden reduction of 10 − 30% continues to be expected over a period of 30 years. We question the real risks of vaccine disease enhancement occurring when implementing Dengvaxia according to the WHO-SAGE recommendation [45]. Here we make such an assessment.

## 2 Results

In this section, the model is validated with hospital admission data for the province of Chiang Mai, Thailand, supplied by the Ministry of Public Health [16,47]. After model validation and parametrization via data matching, two different scenarios for vaccine implementation are used to investigate the vaccine’s manufacturer claim of a “potential reduction in disease burden by 50% when at least 20% of a target population is vaccinated” [23]. Given the prediction horizon of our deterministic system (approximately 10 years) [14, 18], disease outcome dynamics are evaluated for a period of 5 years after implementing vaccination.

In the first scenario, the newly existing imperfect dengue vaccine, Dengvaxia, is implemented in a population without any immunological-profile screening prior to vaccination, where both seronegative and seropositive individuals, from 9 to 45 years of age, are vaccinated. In the second scenario, after population serological screening prior to vaccination, only seropositive individuals are eligible to receive the vaccine.

### 2.1 Model validation

An analysis of real world empirical dengue data reveals that large disease outbreaks are generally followed by smaller epidemics. The explanation for this dynamic is that during a large outbreak the number of susceptibles in the population is reduced, needing time to build up again before another large outbreak occurs. For Chiang Mai, for example, a large outbreak occurred in 1987 followed by some smaller outbreaks, and then a large outbreak in 1994. This outbreak was followed by 15 years of lower disease incidence. Another large outbreak is observed in 2010 with a similar amplitude to that of 1994. While in 2013, the largest outbreak occurred in a 35 year period. However, baseline dengue transmission in susceptible children assures that small epidemics occur continuously in between large outbreaks.

Chaotic behavior exists in many natural systems. In the case of our deterministic system, whose behavior can in principle be predicted for approximately 10 years [13,14,18], Lyapunov exponents calculations were performed in order to quantify its prediction horizon and beyond unpredictability. By using the same parameter set, short periods of the time series simulation could be compared from time to time, with the empirical data. We use the available hospitalized admission data for Chiang Mai presented in Fig. 6 (a). Similarly as found in [18], and as an example of the many different matching possibilities, Fig. 6 (b) and Fig. 6 (c) describe a 5 year period of empirical data, from 2002 to 2007 and from 2011 to May 2016, respectively, where there is good qualitative (and even quantitative) agreement with our model simulation. Note that different patterns of the data were matched with different transients of the model simulation but always using the same parameter set. That is possible given that multiple patterns can occur in the same chaotic attractor of the system. For more information on qualitative features of chaotic systems, see e.g. [46].

In this study, our model is parameterized by matching the last 5 years of available empirical hospitalized dengue data for Chiang Mai with model simulations. In order to classify the dynamic pattern of the model, we discard long transients which would carry information about the initial conditions. The model parameters are fixed as shown in Table 1.

**Table 1.** Estimated parameter set via data matching. The population of Chiang Mai Province in Thailand is *N* = 1.65 million inhabitants [47] and for 2016, when vaccinaton is implemented, the seroprevalence in the age group eligible to receive the vaccine, 9 −45 years old, is estimated to be composed of 15.6% of seronegatives (S) and 13.8% of monotypic immunes (*S*_1_ + *S*_2_).

| Par. | Description | Values | Ref |
| --- | --- | --- | --- |
| $N$ | population size | 1.65 million inhabitants | [47] |
| $\mu$ | birth and death rate | $1/65y$ | [48] |
| $\gamma$ | recovery rate | $52y^{-1}$ | [14, 49] |
| $\beta$ | effective infection rate | $2\gamma$ | [7, 14] |
| $\eta$ | degree of seasonality | 0.35 | [11, 14] |
| $\varphi$ | phase of seasonality | $3/12y$ | [14] |
| $\rho$ | import parameter | $10^{-8}$ | [11, 14] |
| $\alpha$ | temporary cross-immunity rate | $2y^{-1}$ | [14, 50] |
| $\phi = \phi_v$ | ratio of secondary infections<br>contributing to force of infection | 0.9 | [5, 14, 17, 51] |
| $\psi$ | vaccine-disease enhancement factor | 1.1 | [25] |
| $v$ | vaccination rate | $1/1y$ | [20, 21, 24] |
| $\epsilon_p = \epsilon_s$ | proportion of vaccinated population | 4% or 20% per year | [23] |
| $k_1$ | estimated vaccine efficacy<br>for hospitalization under 9y | -0.536 | [24, 25, 38] |
| $k_2$ | estimated vaccine efficacy<br>for hospitalization older 9y | 0.9 | [24, 25, 38] |

**Figure 6.**
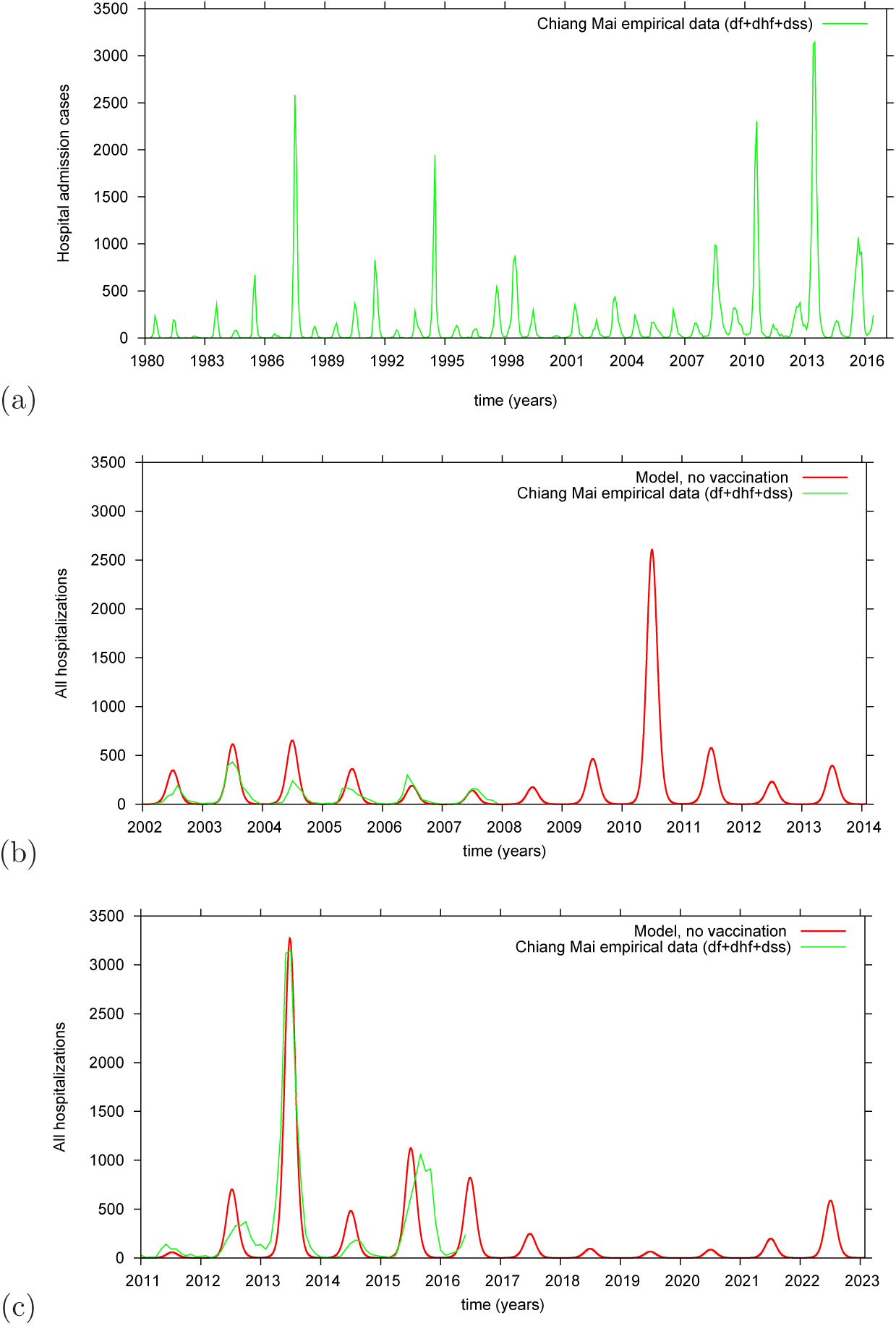
Model validation and parametrization via data matching: (a) reported hospital admission cases in the Chiang Mai Province, Thailand, 1980 to May 2016, (b) real data (in green) compared with model predicted cases without vaccination (in red), from 2002 to 2007 and (c) comparisons for 2011 to May 2016.

### 2.2 Time series analysis for vaccine implementation

In this section, different vaccination strategies are evaluated, 4% and 20% of 9 − 45 year-olds receive vaccine every year.

#### 2.2.1 Impact of imperfect vaccine without immunological screening

In this scenario, from 2016, Dengvaxia is given yearly to 9 − 45 year-olds in Chiang Mai. Both monotypic immune (13.8%) and seronegative (15.6%) individuals are eligible to receive the vaccine without immunological screening. Vaccine efficacies for seropositive and seronegative individuals are as presented in section 1.1.2 and in Table 1.

Figure 7 shows model simulation in the absence of vaccination for the whole population up to 65 years of age (red line). Hospitalizations occur during natural secondary dengue infections. Although children under 9 years and adults older than 45 years are not vaccinated, if susceptible, they can be naturally infected and are able to transmit the virus. Figure 7 also shows hospitalizations predicted by the model when 4% and 20% of age group 9 – 45 years are vaccinated annually during the period 2016 − 2021. As opposed to the claim by Sanofi Pasteur’s modelling group of 50% reduction of cases in 5 years [23], when 4% of the target population is vaccinated (see Fig.7 (a)) total hospitalizations increase by 35% in year 4 after vaccine introduction. Five years after vaccine implementation, hospitalizations are predicted to be more than 70% higher than without vaccine. Thus, increasing vaccination to 20% of the target population, seronegatives and seropositives, per year yields a huge increase in hospitalization (see Fig.7 (b)).

**Figure 7.**
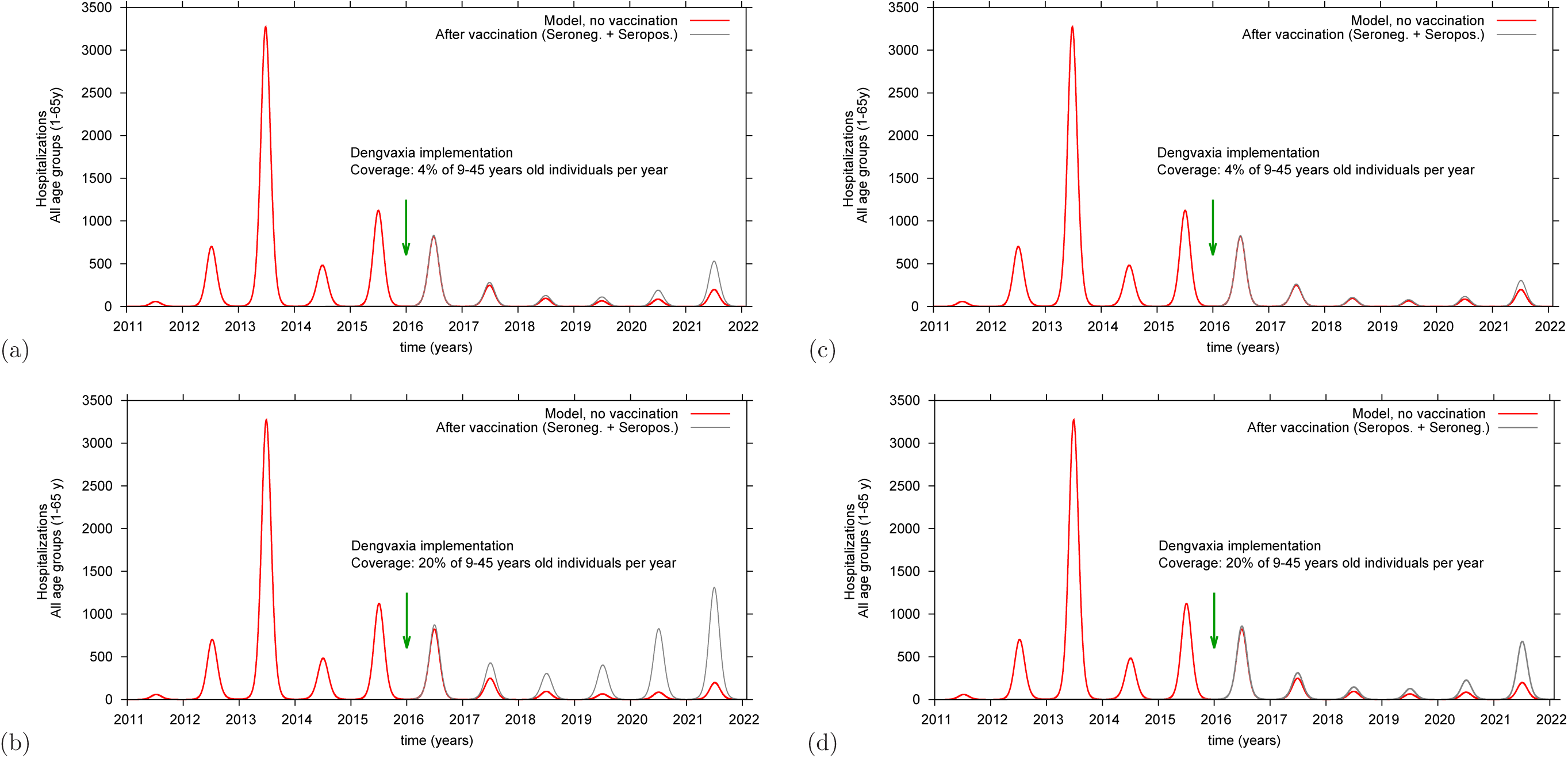
Model simulation of hospitalizations in 1 – 65 year-old individuals for 2016 – 2021. Hospitalizations in the non-vaccinated population is shown with a red line while hospitalizations occurring after yearly vaccination of both seropositives and seronegatives, ages 9 – 45 years, is shown as a grey line. For *ψ* = 1.1, in (a) vaccination coverage is 4% of the population and in (b) vaccination coverage is 20%. For *ψ* = 1.0, vaccination coverage is 4% (c) and in (d) vaccination coverage is 20%.

The parameter *ψ* acts (conservatively) to increase by 10% (*ψ* = 1.1) the risk that vaccinated seronegative individuals who acquire a natural first dengue infection will be hospitalized compared with the risk of hospitalization in unvaccinated monotypic immune 9 – 45 year-olds during a second dengue infection (Fig. 7 (a-b)). The introduction of *ψ* is based on published information showing that the risk of hospitalization of vaccinated seronegatives is increased, presumably expressing higher viremia titers [25, 27]. Even if the risk of hospitalization of dengue infected vaccinated seronegatives and monotypic immunes is the same (*ψ* = 1.0), similar increases in hospitalizations are observed after introduction of vaccine (Fig. 7 (c-d)). At present no data are available to support an assumption of (*ψ* < 1.0).

#### 2.2.2 Impact of serological screening and vaccination of seropositives

A prior assessment of the results of the CYD-TDV clinical trials concluded that vaccination of seropositives resulted in high rates of protective immunity [24, 25, 27] The outcome of screening 4 or 20% of individuals ages 9 − 45 years to identify and then vaccinate only dengue immunes annually is modeled in Fig. 8 and Fig 9. Seronegative individuals are not eligible to receive vaccine. All susceptibles, ages 1 − 8 and 46 − 65 years plus seronegatives, ages 9 − 45 years, may become infected and transmit virus. The majority of these infections will be mild or unapparent. When 4% of seropositives are vaccinated there is a decrease in hospitalizations over the period of 5 years of more than 40%, compared with no vaccination (see blue line in Fig. 8). Yearly vaccination directed at 20% of seropositives, age 9 − 45 will decrease hospitalizations by 70% already in the first two years after vaccination, confining dengue transmission to very low numbers in the following years. (see blues line in Fig. 9).

**Figure 8.**
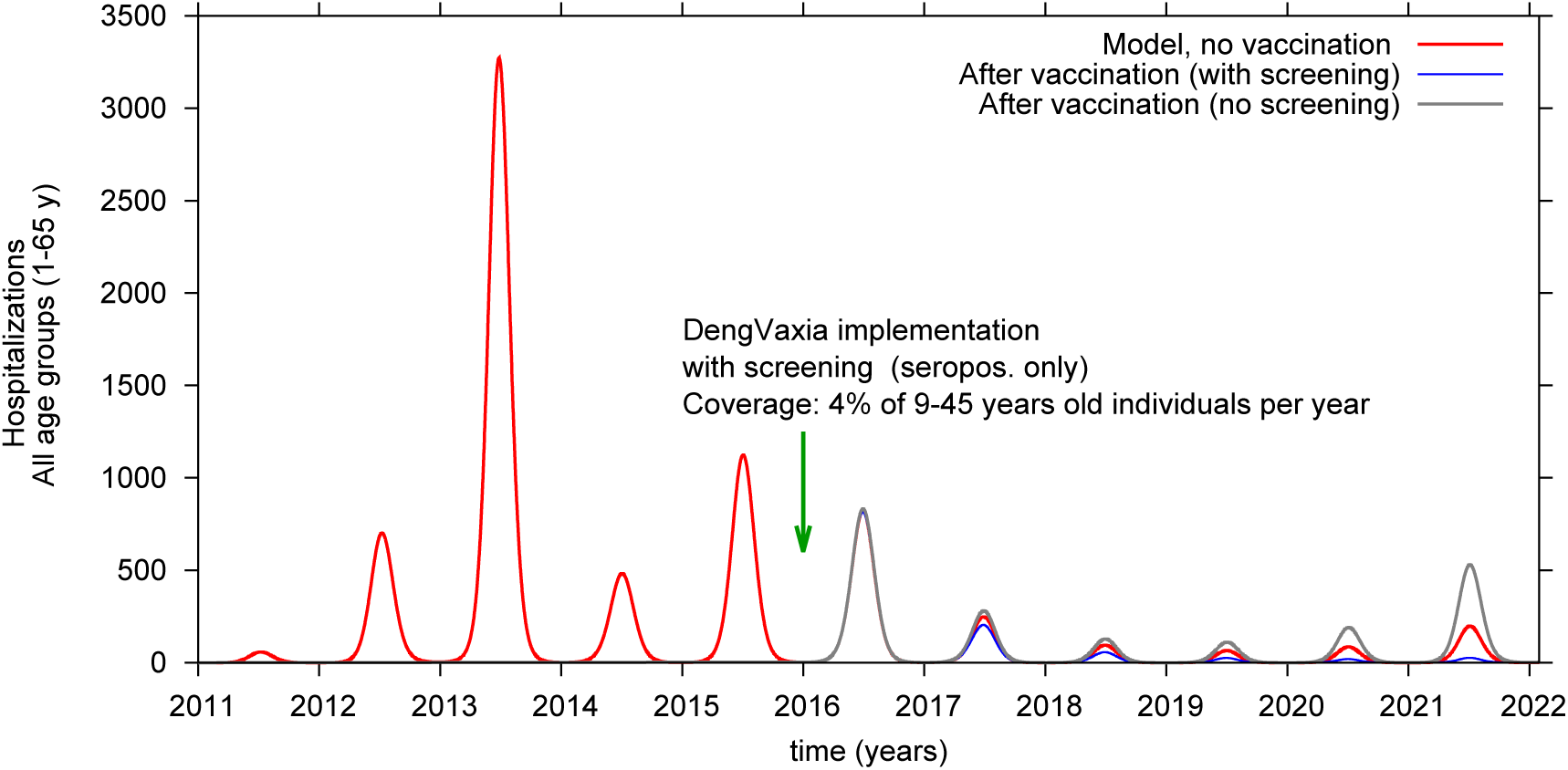
Model simulation of hospitalizations in individuals 1 − 65 years-old implemented in 2016. Only seropositive individuals between 9 − 45 years of age are vaccinated. Yearly vaccine coverage is 4% of 9 − 45 year-olds. Red line is hospitalizations without vaccine, blue line is hospitalizations in vaccinated seropositives who had been serologically screened. Grey line shows hospitalized cases after vaccination of both seropositive and seronegatives.

**Figure 9.**
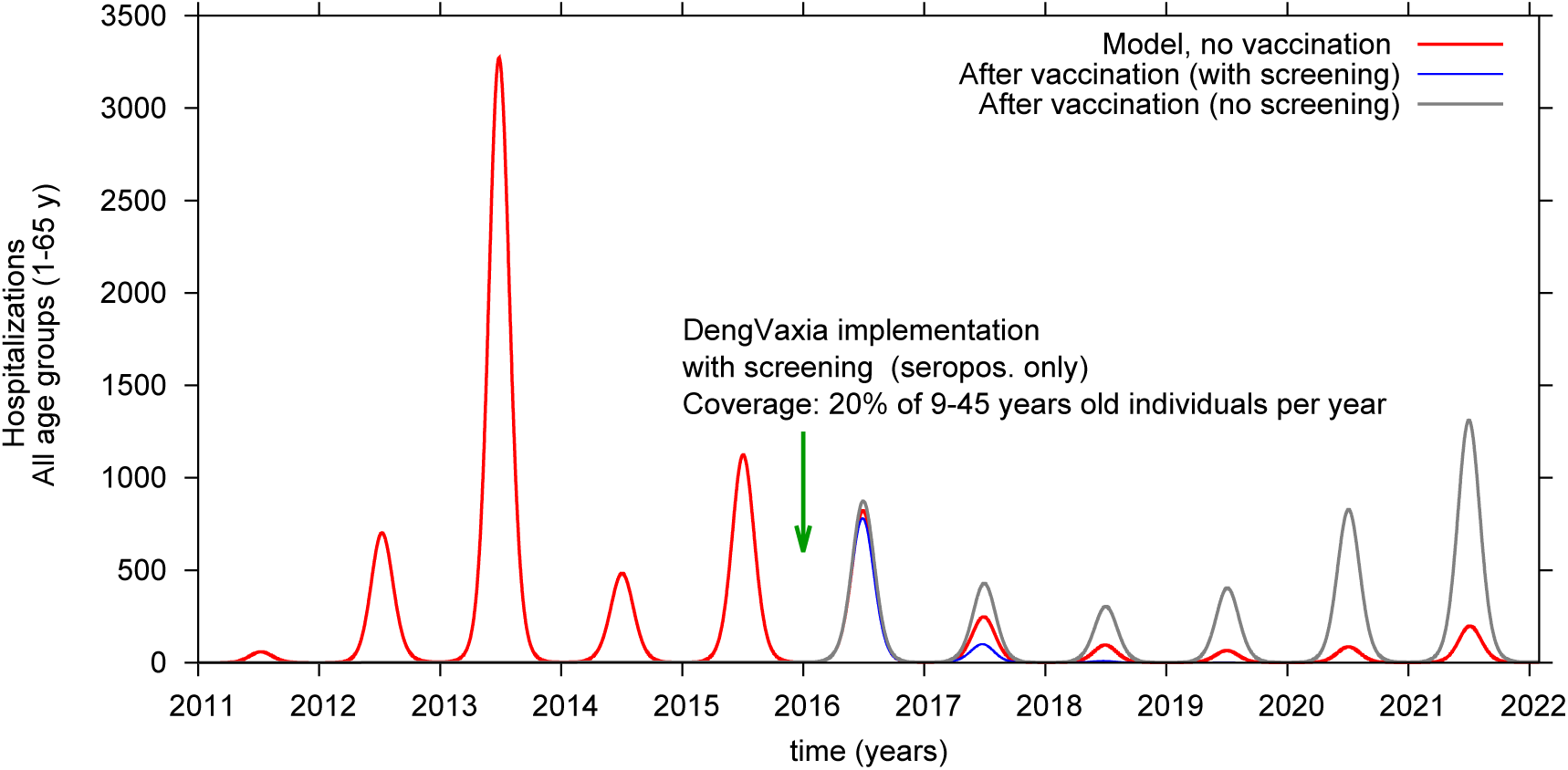
Model simulation of hospitalizations in individuals 1 − 65 years-old implemented in 2016. Only seropositive individuals between 9 − 45 years of age are vaccinated. Yearly vaccine coverage is 20% of 9 − 45 year-olds. Red line is hospitalizations without vaccine, blue line is hospitalizations in vaccinated seropositives who had been serologically screened. Grey line shows hospitalized cases after vaccination of both seropositive and seronegatives.

#### 2.2.3 Impact of vaccination campaigns on hospitalizations of children, ages 1 − 8 years

Fig. 10 plots the impact of vaccinating 9 − 45 year-olds on hospitalizations of children less than 9 years of age. By vaccinating 4% of the seronegative and seropositive target population every year, hospitalizations in children under 9 during the first 3 years stay in the same range as among unvaccinateds (see Fig. 10 (a)). In years 4 and 5 hospitalizations (grey line) increase slightly. When 20% of the target population are vaccinated hospitalizations in children under 9 years increase immediately (see Fig. 10 (b)), year by year. Similar results are also observed for the non-vaccinated group of individuals older than 45 years of age (not shown). However, when 4% of seropositives only, ages 9 − 45 years, are vaccinated yearly there is a gradual reduction in the number of hospitalizations among non-vaccinated young children peaking at 60% five years after vaccine introduction (dark brown line in Fig. 10 (a)). After implementing yearly vaccination of 20% of screened seropositives, there is a major reduction in hospitalizations in unvaccinated children (dark brown line in Fig. 10 (b)), as well in the unvaccinated group of individuals older than 45 years of age (not shown).

**Figure 10.**
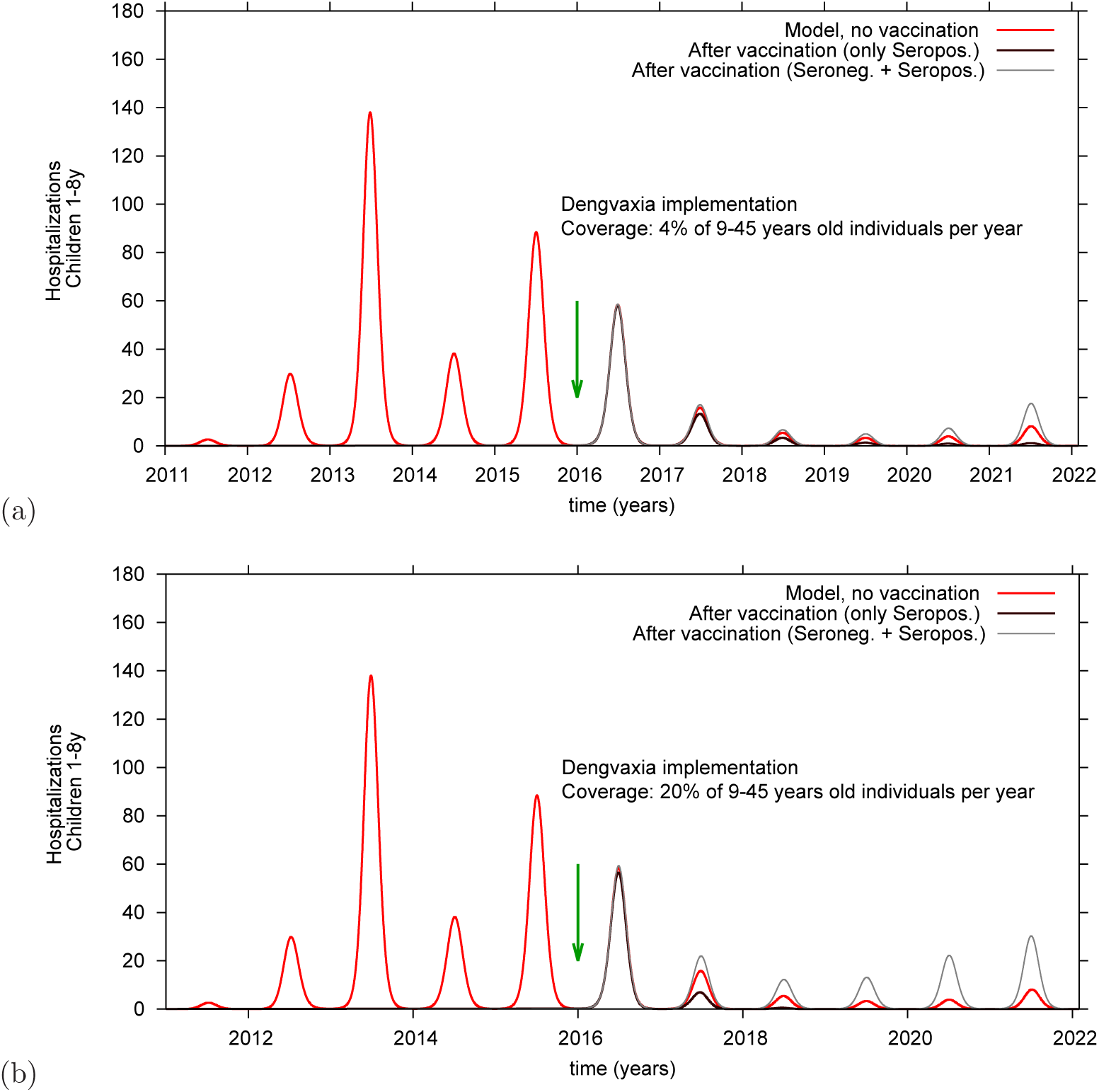
Hospitalizations after implementing yearly vaccination of 4% (a) or 20% (b) of 9 − 45 year-olds. Children under 9 years (and adults older 45 years) are not eligible to receive vaccine. The red line plots hospitalizations, in the absence of vaccine, of 1 − 8 year-olds children, brown line plots hospitalizations after vaccination of seropositives only, showing the impact on non-vaccinated 1 − 8 year-olds. The grey line plots hospitalizations of children under 9 after vaccination of both seropositives and seronegatives, between 9-45 years of age.

## Discussion

The basic purpose of our model is to assess the individual impact and public health implications of the administration of Dengvaxia to seronegative individuals of any age. It has been reported that as of 2015 there were 295 hospitalizations of children receiving Dengvaxia [27]. It is not known if these cases occurred in vaccinated seronegative or seropositive children. However, as nearly 100% of seronegative children receiving vaccine raised dengue antibodies but nonetheless were poorly protected. It was concluded that hospitalizations comprise strong evidence of vaccine enhanced disease [20,21,25]. This assumption is underscored by the observation that vaccinated 2−5 year-old children had a relative risk of hospitalization of 7.45 and six years after being vaccinated, 2.8% of children who had been vaccinated at ages 2 − 5 years were hospitalized [24, 27].

In an early model of vaccination of Asian children using seroprevalence and dengue virus infection data supplied by the manufacturer, it was calculated that vaccination of 2 − 5 year-old children resulted in a 3.5 times higher risk of being hospitalized during a first dengue infection than did unvaccinated monotypic dengue-immune children experiencing a second dengue infection [25]. As the intrinsic risk to experience the dengue vascular permeabilty syndrome decreases rapidly with age, for this model we have chosen a conservative increase in risk of hospitalization of 0.1% during the first dengue infections of vaccinated seronegatives between 9 − 45 years of age. Correspondingly, because at year 2 there was published evidence of 80% protection of vaccinated seropositive children against symptomatic disease accompanying breakthrough dengue infections our model assumes a high rate (90%) of protection after administering dengue vaccine to seropositive individuals [24, 27].

To validate our hypotheses prospective model simulations are compared to hospitalization data from Chiang Mai Province, Thailand. Several of the authors have made extensive use of 35 years of dengue hospitalization data from this Province in earlier epidemiological simulations [13–16, 18].

According to Sanofi, Dengvaxia, given to 20% of individuals 9 − 45 years old, has the potential to reduce dengue disease burden by about 50% within 5 years, achieving a WHO dengue prevention goal and achieving a significant public health impact [23]. On April 15, 2016, SAGE, summarizing work of eight modeling groups commented that “All models predicted that routine vaccination of 9 year-olds with CYD-TDV at 80% vaccine coverage would cause an overall reduction in dengue disease in moderate to high transmission intensity settings (seroprevalence -SP -at age 9 ≥ 50%). This range of transmission intensity covers all the sites selected for the phase 3 trials of CYD-TDV. The impact of vaccination was greatest in high transmission intensity settings (SP9 ≥ 70%), where the reduction in dengue-related hospitalizations predicted by the models ranged from 10% to 30%” over a period of 30 years [26, 27]. However, since seroprevalence is not a uniform variable, a prediction period as long as 30 years raises questions of the validity of this recommendation [45]. It should be noted that when a target population is composed of 70% of partial dengue-immune individuals, the remaining 30% of the population are seronegative.

In this paper, vaccine efficacy in preventing hospitalized dengue cases was estimated using a Bayesian approach [38,40]. Using Sanofis recommendation to vaccinate persons age 9 − 45 years in dengue endemic countries with a high seroprevalence of dengue antibodies, an age-group-structured model was developed to evaluate the impact of vaccination campaigns with and without serological screening for past dengue infections. With this model the impact on hospitalization was assessed when 4 or 20% of individuals ages 9 − 45 years were given vaccine yearly. Our results show that 4% vaccination of unscreened seronegative plus seropositive individuals in the target population, 9−45 years-old, results in a more than 70% increase in hospitalizations in the whole population, 1 − 65 years, during the next 5 years. When vaccine is given to 20% of the target population per year there is an even larger increase in hospitalization, a true public health disaster. When the target population of seronegative and seropositives were vaccinated there was also a negative impact on hospitalization of children under 9 and adults older than 45 years. At 4% vaccination, there was a modest increase in hospitalization but at 20% vaccination, there was a sharp and significant increase in cases in unvaccinated children 1 − 8 years. Overall, our conclusion is that giving Dengvaxia without screening results in a significantly increased disease burden.

However, if vaccination is directed at seropositive individuals, 9 − 45 years of age, a considerable reduction in disease burden is observed. As shown in section 2.2.2, serological screening prior to vaccination resulted in a significant reduction of hospitalized cases in the whole population reducing hospitalization of non-vaccinated individuals, presumably by decreasing the force of infection in the system. Here, when 4% of seropositives, ages 9 − 45 years are vaccinated, a decrease is observed in hospitalizations over the period of 5 years of greater than 40%. Yearly vaccination of 20% of seropositives in the target population results in a decrease in total hospitalizations of 60% within two years after start (cumulative 60% coverage), severely reducing dengue hospitalizations thereafter.

A question that needs to be addressed at all prospective vaccination sites is which screening test to choose to identify dengue-immunes. ELISA IgG tests vary in specificity. Populations in Asia and the Americas will differ in exposure to non-dengue flaviviruses or vaccines. Careful attention to test development and to the specific requirements imposed by varying viral ecologies must be incorporated into program planning. Also, heterogeneities in dengue endemicity may result in marked variation in the prevalence of dengue antibodies in 9 year-old children. As the prevalence of seronegatives increases also there is a risk of an increased disease burden after implementing vaccination campaigns. There have already been objections raised concerning the cost of serological screening [52, 53]. We understand that adding serological testing is a unique requirement in the history of vaccination. However, the increases in disease burden predicted by our model, not only will result in individual harm but also represents a major cost to families and to society. It is this additional cost imposed by the unnecessary burden of vaccine enhanced disease that justifies the allocation of resources to use this “imperfect” vaccine most efficiently and to optimize a beneficial outcome.

## Acknowledgments

Dengue surveillance data were provided by the Bureau of Epidemiology, Department of Disease Control, Ministry of Public Health, Thailand.

